# Less neutralization evasion of SARS-CoV-2 BA.2.86 than XBB sublineages and CH.1.1

**DOI:** 10.1101/2023.09.10.557047

**Authors:** Yanping Hu, Jing Zou, Chaitanya Kurhade, Xiangxue Deng, Hope C. Chang, Debora K. Kim, Pei-Yong Shi, Ping Ren, Xuping Xie

## Abstract

The highly mutated BA.2.86, with over 30 spike protein mutations in comparison to Omicron BA.2 and XBB.1.5 variants, has raised concerns about its potential to evade COVID-19 vaccination or prior SARS-CoV-2 infection-elicited immunity. In this study, we employ a live SARS-CoV-2 neutralization assay to compare the neutralization evasion ability of BA.2.86 with other emerged SARS-CoV-2 subvariants, including BA.2-derived CH.1.1, Delta-Omicron recombinant XBC.1.6, and XBB descendants XBB.1.5, XBB.1.16, XBB.2.3, EG.5.1 and FL.1.5.1. Our results show that BA.2.86 is less neutralization evasive than XBB sublineages. Among all the tested variants, CH.1.1 exhibits the greatest neutralization evasion. In comparison to XBB.1.5, the more recent XBB descendants, particularly EG.5.1 and FL.1.5.1, display increased resistance to neutralization induced by parental COVID-19 mRNA vaccine and a BA.5-Bivalent-booster. In contrast, XBC.1.6 shows a slight reduction but remains comparable sensitivity to neutralization when compared to BA.5. Furthermore, a recent XBB.1.5-breakthrough infection significantly enhances the breadth and potency of cross-neutralization. These findings reinforce the expectation that the upcoming XBB.1.5 mRNA vaccine would likely boost the neutralization of currently circulating variants, while also underscoring the critical importance of ongoing surveillance to monitor the evolution and immune evasion potential of SARS-CoV-2 variants.

## Main text

Severe Acute Respiratory Syndrome Coronavirus 2 (SARS-CoV-2), the causative agent of the COVID-19 pandemic, continues evolving and diverging regionally and globally. Since its initial outbreak in December 2019^1^, SARS-CoV-2 has constantly acquired genetic changes for improved fitness and immune evasion. Such adaptations have given rise to variants including Alpha, Beta, Delta, and Omicron, each of which has caused surges in infections worldwide^2^. The emergence of Omicron represents a significant shift in the evolution trajectory of SARS-CoV-2. Omicron carries more than 30 amino acid changes in its spike compared to non-Omicron variants, which account for its remarkable transmissibility and immune evasion^3-6^. Since its first detection in December 2021^7^, Omicron has rapidly become dominating and subsequently evolving into a swarm of Omicron subvariants^8^. Several Omicron sublineages, such as BA.2, BA.5, and XBB, have caused waves of infections globally. Among these sublineages, the XBB variant, a recombination of BA.2.10 and BA.2.75 sublineages, has displayed significant immune evasion^9-14^ and rapidly outcompeted the previously dominant BA.5 variant since early 2023. As a result, the Food and Drug Administration (FDA) recommends the use of a monovalent XBB sublineage as the vaccine composition for the coming year 2023-2024^15^. Currently, several other descendants of XBB, including XBB.1.16, XBB.2.3, EG.5, and FL.1.5.1 (Fig.S1), are now overriding the XBB.1.5 dominance on a global scale (https://covid.cdc.gov/covid-data-tracker).

More recently, a subvariant derived from BA.2, known as BA.2.86, has raised concerns about its potential to evade immunity induced by COVID-19 vaccination or prior natural infections. BA.2.86 carries over 30 amino acid changes in its spike when compared to both BA.2 and XBB.1.5, a similar magnitude of amino acid variation as previously observed between the BA.1 and Delta variants. The World Health Organization designated it as a variant under monitoring on 17 August 2023. As of 5 September 2023, there have been 41 reported sequences of BA.2.86 from 12 countries in the GISAID database, which is likely underestimated due to limited sampling. Additionally, two other circulating non-XBB Omicron subvariants CH.1.1 (a descendant of BA.2.75 sublineage) and XBC.1.6 (a recombination of Delta and Omicron sublineage BA.2) have also drawn attention due to their great immune evasion capabilities^13,16,17^. As of 29 August 2023, CH.1.1 has expanded from Southeast Asia to over 86 countries, while XBC.1.6 subvariant has steadily increased in prevalence in the Philippines and Australia (https://outbreak.info/situation-reports). Therefore, it is of utmost importance to compare the ability of these circulating variants to evade immunity generated by vaccination or previous natural infections.

The goal of this study was to assess the neutralization of human sera after vaccination and/or prior SARS-CoV-2 infections against these newly emerged Omicron sublineages (XBB.2.3, XBB.1.16, EG.5.1, FL.1.5.1, CH.1.1, XBC.1.6, and BA.2.86). To enable accurate neutralization measurement, we engineered the complete *spike* gene from each of these Omicron sublineages into the backbone of mNeonGreen (mNG) reporter USA-WA1/2020 SARS-CoV-2 (Fig.S2). Compared with wild-type USA-WA1/2020 (a strain isolated in January 2020), the insertion of the *mNG* gene at open-reading-frame-7 of the viral genome attenuated the virus *in vivo*^18^. Thus, the engineered live-attenuated mNG viruses can be used safely in a BSL3 facility with the correct procedures for neutralization^19^. The recovered recombinant viruses demonstrated high infectivity on VeroE6 cells with titers exceeding 10^7^ PFU/ml. Before their use in determining the 50% fluorescent focus-reduction neutralization titers (FFRNT_50_) of human sera, all recombinant viruses were verified by Sanger sequencing to ensure no undesired mutations in their genomes.

We analyzed four human serum panels with distinct vaccination and/or SARS-CoV-2 infection histories. The first panel comprised 28 sera obtained from individuals 14-32 (median 23) days post BA.5-bivalent-booster (referred to as BA.5-bivalent-booster sera); these sera were collected from September 30 to October 22, 2022 (Table S1). The second panel included 20 sera from individuals who had previously contracted SARS-CoV-2 and subsequently received a BA.5-bivalent-booster 15-31 (median 21) days ago (referred to as BA.5-bivalent-booster-infection sera). Samples collection for this panel took place from October 4 to 22, 2022 (Table S2). The infection history for the second panel was confirmed by positive nucleocapsid antibody detection or SARS-CoV-2 RT-PCR^10^. The third panel comprised 42 sera collected from vaccinated individuals 15-117 days (median 47) after a breakthrough infection with XBB.1.5 variant (referred to as XBB.1.5-infection-parental-mRNA vaccine sera). These samples were obtained between February 2 and May 23, 2023 (Table S3). The fourth panel included 19 sera collected 17-103 days (median 42) after XBB.1.5-breakthrough infection in individuals who had previously received a BA.5-bivalent booster (referred to as XBB.1.5-infection-BA.5-bivalent-plus-parental-mRNA-vaccine sera). These samples were collected between February 16 and May 18, 2023 (Table S4). For the third and fourth panels, the XBB.1.5 infection was confirmed through the sequencing of nasopharyngeal specimens; however, the infection history prior to the XBB.1.5-infection could not be determined. It is important to note that individuals in all four panels had received 2-4 doses of the parental monovalent mRNA vaccine before receiving the BA.5-bivalent-booster or experiencing the XBB.1.5-infection. Tables S1-4 summarize the serum information and neutralization titers for each serum panel.

BA.5-bivalent-booster sera neutralized USA-WA1/2020-, BA.5-, XBB.1.5-, XBB.1.16-, XBB.2.3-, EG.5.1-, FL.1.5.1-, CH.1.1-, XBC.1.6-, and BA.2.86-spike mNG SARS-CoV-2 with geometric mean titers (GMTs) of 3576, 290, 42, 39, 48, 29, 31, 23, 152 and 67, respectively (Figure 1A and Table S1). The GMTs against BA.5-, XBB.1.5-, XBB.1.16-, XBB.2.3-, EG.5.1-, FL.1.5.1-, CH.1.1-, XBC.1.6- and BA.2.86-spike viruses were 12.3, 85.1, 91.7, 74.5, 123.3, 115.4, 155.5, 23.5 and 53.4-fold lower than the GMT against the USA-WA1/2020, respectively (Figure 1A). Compared with the GMT against the BA.5-spike, the GMTs against XBB.1.5-, XBB.1.16-, XBB.2.3-, EG.5.1-, FL.1.5.1-, CH.1.1-, XBC.1.6- and BA.2.86-spike were reduced by 6.9, 7.4, 6, 10, 9.4, 12.6, 1.9, and 4.3-fold, respectively.

**Figure 1.**
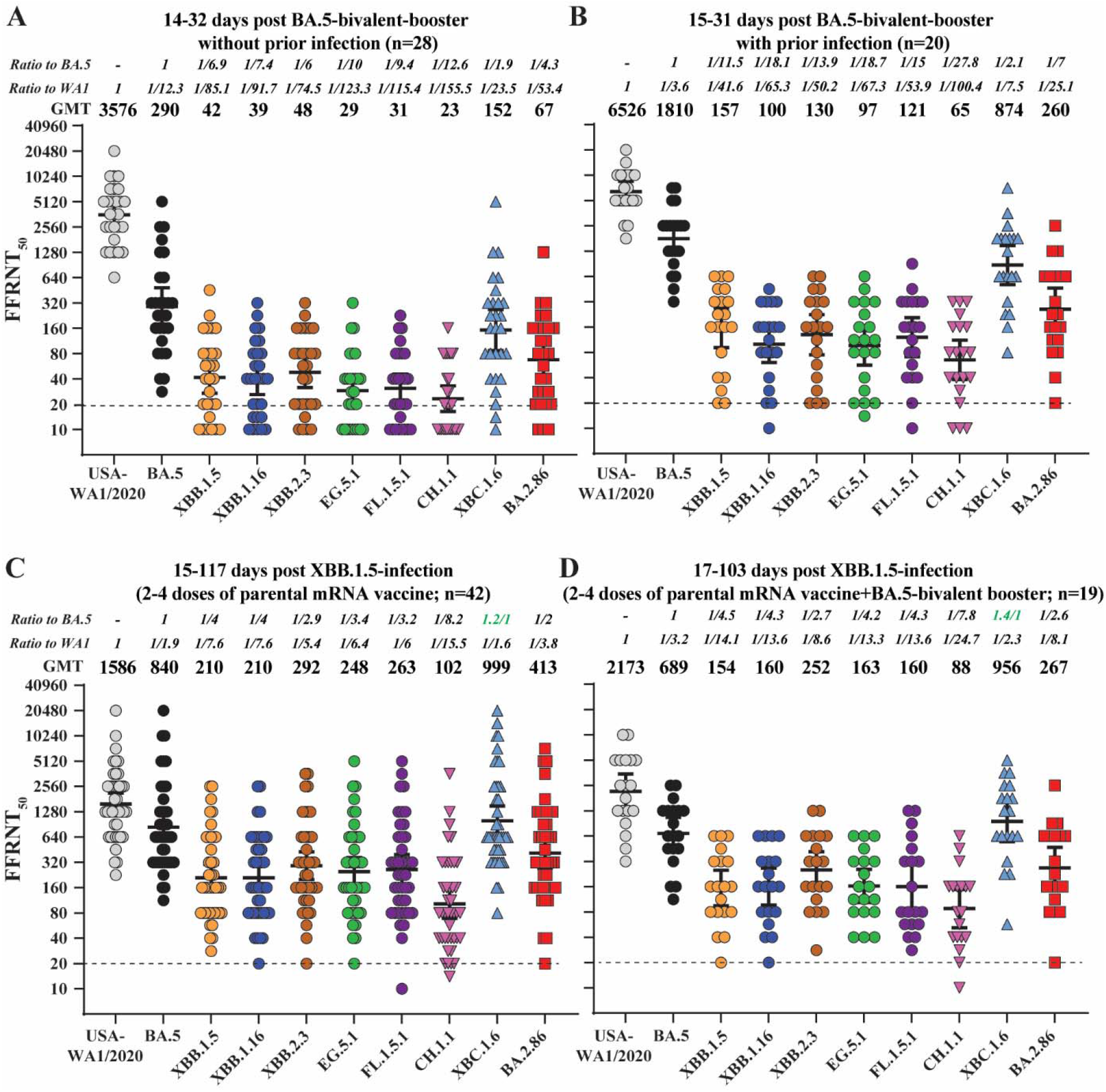
Neutralization titers against Omicron sublineages. (A) FFRNT_50_ of 28 sera collected after BA.5-bivalent booster from individuals without prior SARS-CoV-2 infection. (B) FFRNT_50_ of 20 sera collected after BA.5-bivalent-booster from individuals with prior SARS-CoV-2 infection. (C) FFRNT_50_ of 42 sera collected after XBB.1.5-breakthrough infection from individuals with parental mRNA vaccination. (D) FFRNT_50_ of 19 sera collected after XBB.1.5-breakthrough infection from individuals with parental mRNA vaccination plus BA.5-bivalent booster. The solid lines and numeric values above each panel indicate the geometric mean titers (GMTs). The error bars represent the 95% confidence intervals (Cis). The fold reduction in GMT against each Omicron sublineage, compared with the GMT against USA-WA1/2020 or BA.5-spike, is shown in italic font. The green italic numbers indicate more sensitive to neutralization. The dotted line indicates the limit of detection of FFRNT_50_. FFRNT_50_ of <20 was treated as 10 for plotting purposes and statistical analysis. The *p* values (determined using the Wilcoxon matched-pairs signed-rank test) for group comparison of GMTs are shown in Table S5.

BA.5-bivalent-booster-infection sera neutralized USA-WA1/2020-, BA.5-, XBB.1.5-, XBB.1.16-, XBB.2.3-, EG.5.1-, FL.1.5.1-, CH.1.1-, XBC.1.6- and BA.2.86-spike mNG SARS-CoV-2 with GMTs of 6526, 1810, 157, 100, 130, 97, 121, 65, 874 and 260, respectively (Figure 1B and Table S2). The GMTs against BA.5-, XBB.1.5-, XBB.1.16-, XBB.2.3-, EG.5.1-, FL.1.5.1-, CH.1.1-, XBC.1.6- and BA.2.86-spike were 3.6, 41.6, 65.3, 50.2, 67.3, 53.9, 100.4, 7.5 and 25.1-fold lower than the GMT against the USA-WA1/2020, respectively (Figure 1B). Compared with the GMT against the BA.5-spike, the GMTs against XBB.1.5-, XBB.1.16-, XBB.2.3-, EG.5.1-, FL.1.5.1-, CH.1.1-, XBC.1.6-, and BA.2.86-spike were reduced by 11.5, 18.1, 13.9, 18.7, 15, 27.8, 2.1 and 7-fold, respectively. Compared with BA.5-bivalent-booster sera without infection history, BA.5-bivalent-booster-infection sera increased the neutralizing GMTs against USA-WA1/2020, BA.5-, XBB.1.5-, XBB.1.16-, XBB.2.3-, EG.5.1-, FL.1.5.1-, CH.1.1-, XBC.1.6- and BA.2.86-spike by 1.8, 6.2, 3.7, 2.6, 2.7, 3.3, 3.9, 2.8, 5.8 and 3.9-fold, respectively (compare Fig 1B versus 1A).

XBB.1.5-infection-parental-mRNA vaccine sera neutralized USA-WA1/2020-, BA.5-, XBB.1.5-, XBB.1.16-, XBB.2.3-, EG.5.1-, FL.1.5.1-, CH.1.1-, XBC.1.6- and BA.2.86-spike mNG SARS-CoV-2 with GMTs of 1586, 840, 210, 210, 292, 248, 263, 102, 999 and 413, respectively (Figure 1C and Table S3). The GMTs against BA.5-, XBB.1.5-, XBB.1.16-, XBB.2.3-, EG.5.1-, FL.1.5.1-, CH.1.1-, XBC.1.6-, and BA.2.86-spike were 1.9, 7.6, 7.6, 5.4, 6.4, 6.0, 15.5, 1.6 and 3.8-folds lower than the GMT against the USA-WA1/2020, respectively (Figure 1C). Compared with the GMT against the BA.5-spike, the GMTs against XBB.1.5-, XBB.1.16-, XBB.2.3-, EG.5.1-, FL.1.5.1-, CH.1.1-, and BA.2.86-spike were reduced by 4, 4, 2.9, 3.4, 3.2, 8.2, and 2-fold, respectively. However, the GMT against XBC.1.6-spike was slightly higher than the GMT against BA.5-spike.

XBB.1.5-infection-BA.5-bivalent-parental-mRNA-vaccine sera neutralized USA-WA1/2020-, BA.5-, XBB.1.5-, XBB.1.16-, XBB.2.3, EG.5.1-, FL.1.5.1-, CH.1.1, XBC.1.6, and BA.2.86-spike mNG SARS-CoV-2 with GMTs of 2173, 689, 154, 160, 252, 163, 160, 88, 956, and 267, respectively (Figure 1D and Table S4). The GMTs against BA.5-, XBB.1.5-, XBB.1.16-, XBB.2.3, EG.5.1-, FL.1.5.1-, CH.1.1, XBC.1.6, and BA.2.86-spike were 3.2, 14.1, 13.6, 8.6, 13.3, 13.6, 24.7, 2.3, and 8.1-fold lower than the GMT against the USA-WA1/2020-spike SARS-CoV-2, respectively (Figure 1D). Compared with the GMT against the BA.5, the GMTs against XBB.1.5-, XBB.1.16-, XBB.2.3-, EG.5.1-, FL.1.5.1-, CH.1.1- and BA.2.86-spike were reduced by 4.5, 4.3, 2.7, 4.2, 4.3, 7.8 and 2.6-fold, respectively. Interestingly, in comparison to the fourth panel sera, the third panel sera exhibited overall higher neutralizing GMTs against most of variants except XBC.1.6 and USA-WA1/2020 (Compare Fig 1D versus 1C), which may be due to different immune imprinting resulting from vaccinations and/or prior SARS-CoV-2 infections^16^.

Overall, our results indicate that BA.2.85 can evade neutralization elicited by parental and BA.5 bivalent mRNA vaccine or previous SARS-CoV-2 infections. Surprisingly, despite substantial changes in its spike protein, BA.2.85 exhibited lower immune evasion capabilities compared to XBB descendants and CH1.1. Notably, CH.1.1 displayed the greatest immune evasion among all the tested variants, indicating its potential for wider global spread. Conversely, XBC.1.6 showed the least evasion to neutralization. When compared to neutralizing GMTs against BA.5, the neutralizing GMTs against XBC.1.6 were slightly lower (less than 1-fold reduction) in individuals who had received vaccination and a BA.5-bivalent booster (Figure 1A-B) and became higher after XBB.1.5-breaktrhough infection within our cohort (Figure 1C-D). Consistently, XBB descendants showed substantial evasion of neutralization elicited by parental and BA.5-bivalent mRNA vaccine or prior infections^10^. Interestingly, the more recent XBB descendants, particularly EG.5.1 and FL.1.5.1, exhibited more resistance to neutralization induced by vaccination and BA.5-booster or prior SARS-CoV-2 infections when compared to the earlier XBB descendant XBB.1.5. This observation aligns with the epidemiological data from earlier 2023 to the present, reflecting changes in prevalent XBB sublineages. However, when neutralization was boosted by XBB.1.5-breakthrough infection, all XBB descendants showed comparable sensitivity to neutralization. Our results highlight that hybrid immunity, induced by vaccination and prior infections, can enhance the magnitude and breadth of neutralization. More importantly, XBB.1.5-breakthrough infection led to broader and more robust neutralization against the currently circulating Omicron sublineages, regardless of whether a BA.5 bivalent booster was administrated. These results reinforce the expectation that the upcoming XBB.1.5 mRNA vaccine would likely elevate the neutralization against circulating Omicron variants.

The study has several limitations. First, we have not examined the antiviral roles of non-neutralizing antibodies and cell-mediated immunity. These two immune components, together with neutralizing antibodies, protect patients from severe disease and death^20,21^. Unlike neutralizing antibodies, many T cell epitopes after vaccination or natural infection are preserved in Omicron spikes^22^. However, robust antibody neutralization is critical to prevent viral infection^23^. Second, we have not dissected the spike mutations that contribute to the observed immune evasion of the newly emerged Omicron sublineages. It remains elusive how/what spike mutations drive the different immune evasion between CH.1.1 and the two highly mutated BA.2.86 and XBC.1.6 (Figure S2). Third, we do not know the baseline of neutralization titers before vaccination or infection. Fourth, the current results do not allow a direct comparison of neutralization from diverse immune backgrounds or baselines resulting from the vaccine type, number of vaccine doses, history of infection (when and which variants), geographic and/or demographics. Such immune background complex may explain some subtle neutralization differences as reported by other groups^24,25^. Despite these limitations, our laboratory findings would guide vaccine strategy against both current and future Omicron sublineages. Considering the co-circulation of multiple immune-evasive SARS-CoV-2 variants with distinct genetic changes, we must continue vigilantly monitoring the ongoing evolution of SARS-CoV-2 and its potential to further elude existing public health measures.

## Acknowledgments

We thank many laboratories, colleagues, as well as GISAID staffs, who deposited, distributed, and shared the original sequence and analysis data. P.-Y.S. and X.X. were supported by NIH contract HHSN272201600013C, and awards from the Sealy & Smith Foundation, the Kleberg Foundation, the John S. Dunn Foundation, the Amon G. Carter Foundation, the Summerfield Robert Foundation, and Edith and Robert Zinn. We thank the participants from whom the serum specimens were obtained. The funders had no role in study design, data collection, analysis, decision to publish, or preparation of the manuscript.

## Author contributions

Conceptualization, P.R., X.X..; Methodology, J.Z., Y.H., C.K., X.D., H.C.C., D.K., P.R., X.X.; Investigation, J.Z., Y.H., C.K., X.D., H.C., D.K., P.R., X.X.; Resources, P.-Y.S., P.R., X.X.; Data Curation, J.Z., Y.H., C.K., X.D., P.R., X.X.; Writing-Original Draft, J.Z., Y.H., P.R., X.X.; Writing-Review & Editing, J.Z., Y.H., C.K., X.D., H.C., D.K., P.-Y.S., P.R., X.X.; Supervision, P.R., X.X.; Funding Acquisition, P.-Y.S., P.R., X.X..

## Competing interests

P.-Y.S. and X.X. have filed a patent on the SARS-CoV-2 reverse genetic system. Other authors declare no competing interests.

## Methods

### Ethical statement

The research protocol regarding the use of human serum specimens was reviewed and approved by the University of Texas Medical Branch (UTMB) Institutional Review Board (IRB number 20-0070). No informed consent was required since these deidentified sera were leftover specimens from the routine standard of care and diagnostics before being discarded. No diagnosis or treatment was involved either. The use of human serum specimens in this study was reviewed and approved by the University of Texas Medical Branch (UTMB) Institutional Review Board (IRB number 20-0070).

All virus work was performed in a biosafety level 3 (BSL-3) laboratory with redundant fans in the biosafety cabinets at The University of Texas Medical Branch at Galveston. All personnel wore powered air-purifying respirators (Breathe Easy, 3M) with Tyvek suits, aprons, booties, and double gloves.

### Cells

VeroE6 (ATCC® CRL-1586) purchased from the American Type Culture Collection (ATCC, Bethesda, MD) and Vero E6 cells expressing TMPRSS2 (JCRB1819) purchased from SEKISUI XenoTech, LLC were maintained in a high-glucose Dulbecco’s modified Eagle’s medium (DMEM) containing 10% fetal bovine serum (FBS; HyClone Laboratories, South Logan, UT) and 1% penicillin/streptomycin at 37°C with 5% CO_2_. Culture media and antibiotics were purchased from ThermoFisher Scientific (Waltham, MA). Both cell lines were tested Mycoplasma negative.

### Human Serum

Three panels of human sera collected at UTMB were used in the study. Samples were collected based on availability. Varied ages with both genders are included. The population contains varied races or ethnicities, including white, Hispanic, black, and Asian. Subjects have received at least two doses of the COVID-19 vaccine with or without evidence of SARS-CoV-2 infection. The first panel consisted of 28 sera collected from individuals 14-32 (median 23) days after BA.5-bivalent-booster from Pfizer. This panel had been tested negative for SARS-CoV-2 nucleocapsid protein expression using Bio-Plex Pro Human IgG SARS-CoV-2 N/RBD/S1/S2 4-Plex Panel (Bio-rad). The second panel consisted of 20 sera from individuals who were previously infected by SARS-CoV-2 vaccinated with 2-4 doses of parental mRNA vaccine, and received a BA.5-bivalent-booster 15-31 (median 21) days before serum collection. The genotypes of the infecting SARS-CoV-2 variants could not be determined for the second serum panel. The panel 3 and 4 samples were collected 15-117 days after XBB.1.5 infection (as determined by RT-PCR) from individuals who had received 2-4 doses of mRNA vaccines and/or Bivalent booster. However, the infection history of individuals from panel 3 and 4 was not determined. Patient information was completely deidentified from all specimens. No informed consent was required because these deidentified sera were leftover specimens from standard care and diagnostics before being discarded. The use of human sera for this study was reviewed and approved by the UTMB IRB (number 20-0070). The de-identified human sera were heat-inactivated at 56°C for 30 min before the neutralization test. The serum information is presented in Table S1-4.

### Generation of recombinant Omicron sublineages-mNG SARS CoV-2

Recombinant Omicron sublineage XBB.2.3-, XBB.1.16-, EG.5.1-, FL.1.5.1-, CH.1.1-, XBC.1.6- and BA.2.86-spike mNG SARS-CoV-2s was constructed by engineering the complete *spike* gene from the indicated variants into an infectious cDNA clone of mNG USA-WA1/2020 as reported previously^26,27^. The full-length infectious cDNA clone of SARS-CoV-2 was assembled by *in vitro* ligation followed by *in vitro* transcription to synthesize the viral genomic RNA. The full-length RNA transcripts were electroporated in VeroE6-TMPRSS2 cells to recover the viruses.

Viruses were rescued post 2-3 days after electroporation and served as P0 stock. P0 stock was further passaged once on Vero E6 cells to produce P1 stock. The reason for using Vero E6 cells (rather than using Vero E6-TMPRSS2) to prepare the P1 virus is that the infectivity of the P1 virus can be affected by the cell types; since our established FFRNT assay uses VeroE6 cells, we chose to prepare the P1 viruses using the same VeroE6 cells. The *spike* gene was sequenced from all P1 stock viruses to ensure no undesired mutation. The infectious titer of the P1 virus was quantified by fluorescent focus assay on Vero E6 cells. The P1 virus was used for the neutralization test. The protocols for the mutagenesis of mNG SARS-CoV-2 and virus production were reported previously. All virus preparation and neutralization assays were carried out at the biosafety level 3 (BSL-3) facility at the University of Texas Medical Branch at Galveston.

### Fluorescent focus reduction neutralization test (FFRNT)

Neutralization titers of human sera were measured by FFRNT using BA.5-, XBB.2.3-, XBB.1.16-, EG.5.1-, FL.1.5.1-, CH.1.1-, XBC.1.6- and BA.2.86-spike mNG SARS-CoV-2s at BSL-3 using a previous established FFRNT protocol^28^. Briefly, 2.5 × 10^4^ VeroE6 cells per well were seeded in 96-well plates (Greiner Bio-one(tm)). The cells were incubated overnight. On the next day, each serum was 2-fold serially diluted in the culture medium with the first dilution of 1:20 (final dilution range of 1:20 to 1:20,480). The diluted serum was incubated with 100-150 FFUs of mNG SARS-CoV-2 at 37 °C for 1 h, after which the serum virus mixtures were loaded onto the pre-seeded Vero E6 cell monolayer in 96-well plates. After 1 h infection, the inoculum was removed and 100 μl of overlay medium (supplemented with 0.8% methylcellulose) was added to each well. After incubating the plates at 37°C for 16 h, raw images of mNG foci were acquired using Cytation™ 7 (BioTek) armed with 2.5× FL Zeiss objective with a wide-field of view and processed using the Gen 5 software settings (GFP [469,525], threshold 4000 and object selection size 50-1000 μm). The foci in each well were counted using the Gen5 software and normalized to the non-serum-treated controls to calculate the relative infectivities. The FFRNT_50_ value was defined as the minimal serum dilution that suppressed >50% of fluorescent foci. The neutralization titer of each serum was determined in duplicate assays, and the geometric mean was taken. Tables S2-4 summarize the FFRNT_50_ results. Data were initially plotted in GraphPad Prism 9 software and assembled in Adobe Illustrator. FFRNT50 of <20 was treated as 10 for plotting purposes and statistical analysis.

### Statistics & Reproducibility

No statistical method was used to predetermine the sample size. The samples were collected based on availability. No data were excluded from the analyses. The experiments were not randomized. Patient information was blinded in the study. The investigators were blinded to sample identity during data collection and/or analysis. The experiments were performed in duplication. All attempts at replication were successful.

Continuous variables were summarized as the geometric mean with 95% confidence intervals or median. Sera with undetectable (<20) antibody titers were assigned an antibody titer of 10, for purposes of GMT calculations or statistical comparisons. Comparison between neutralization titers was performed using a Wilcoxon matched-pairs signed-rank test using GraphPad Prism 9.0. Absolute *P* values were provided. *P*□<□0.05 was considered statistically significant. Images were assembled using Adobe Illustrator.

## Data availability

The raw data that support the findings of this study are shown in the Source data files. The sequence of SARS-CoV-2 variants can be accessed through GISAID (https://gisaid.org) with the following codes: XBB.1.16 (EPI_ISL_17030006), XBB.2.3 (EPI_ISL_16475206), EG.5.1 (EPI_ISL_17700360), FL.1.5.1 (EPI_ISL_18224090), CH.1.1 (EPI_ISL_16907910), XBC.1.6 (EPI_ISL_18161082) and BA.2.86 (EPI_ISL_18110065). The sequence of SARS-CoV-2 mNG can be found in the supplementary information of our previous study^19^.

