## Supplementary figure S1-2 and table S1-5 for "Less neutralization evasion of SARS-CoV-2 BA.2.86 than XBB sublineages and CH.1.1"

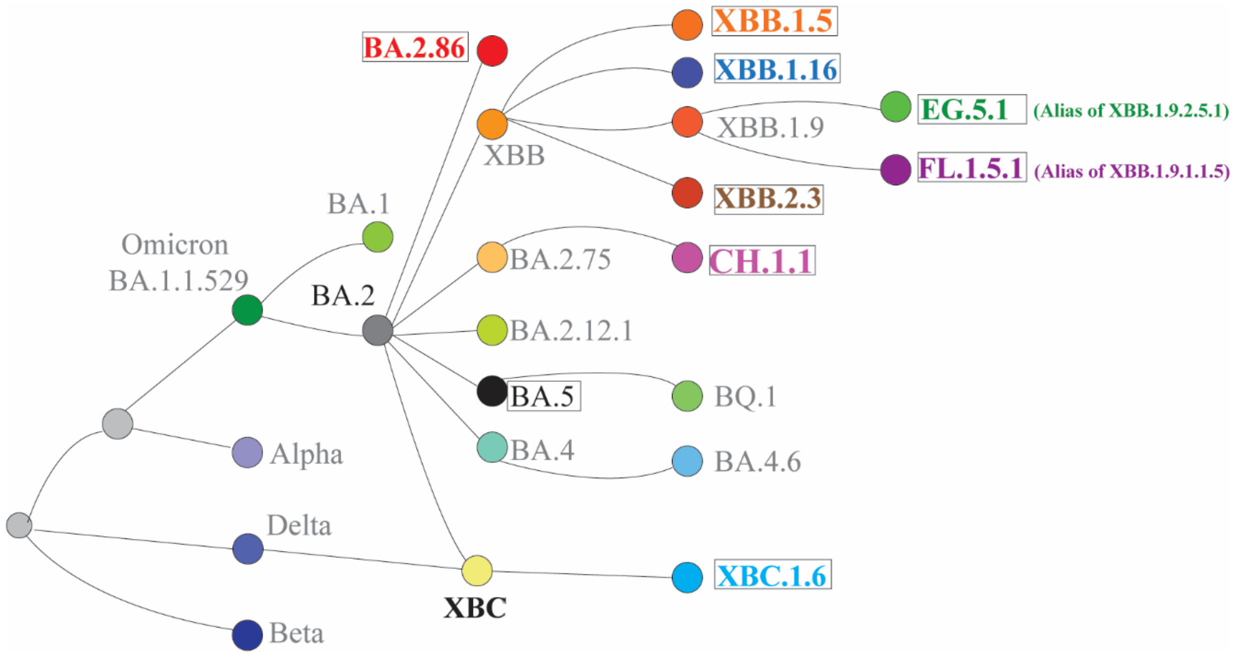


Figure S1. Phylogenetic relationship of SARS-CoV-2 subvariants. The figure was replotted based on the diagram from COVID Data Tracker (https://covid.cdc.gov/covid-data-tracker/#variant-proportions). The subvariants tested in this study were boxed.

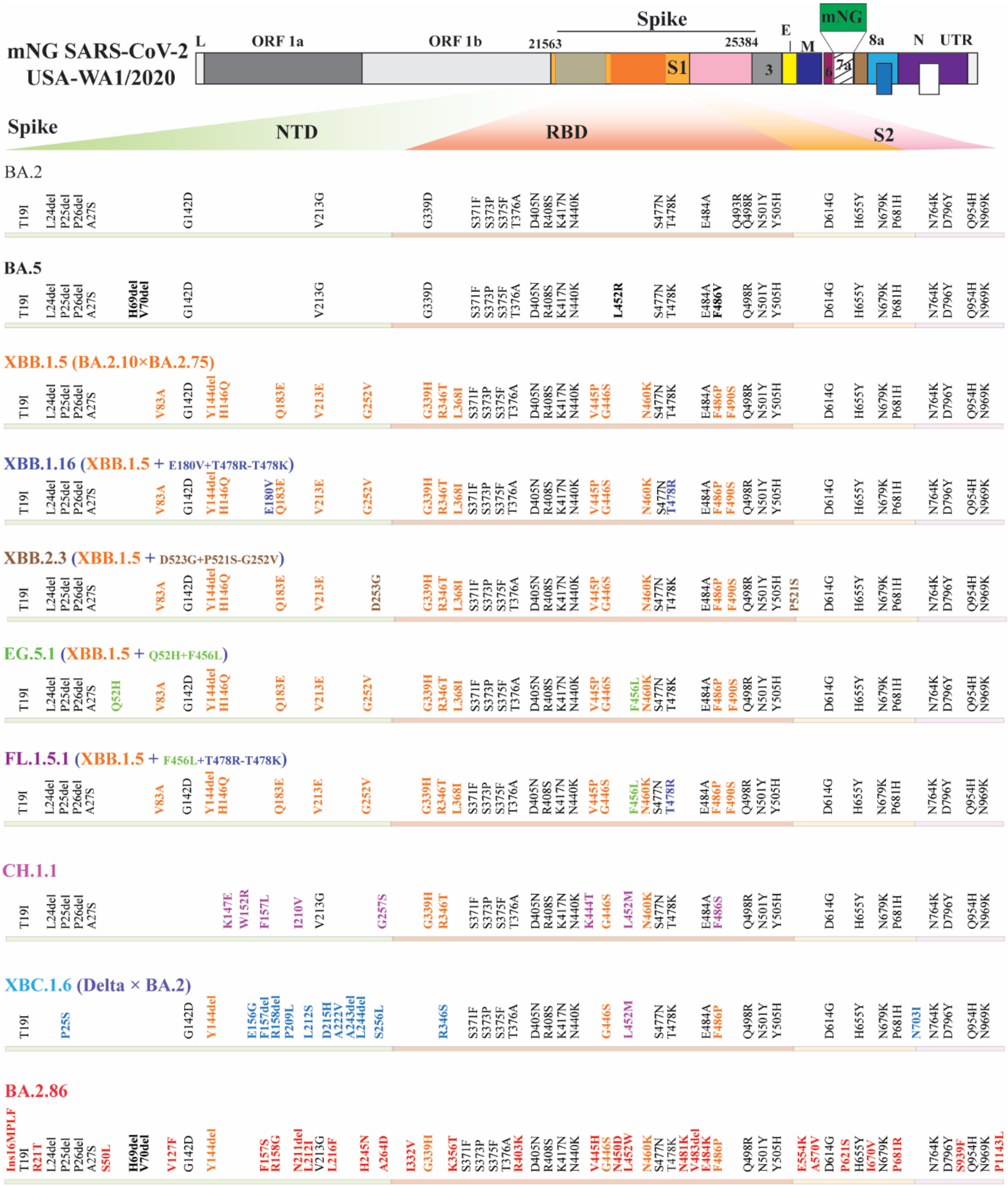


Figure S2. Construction of mNG SARS-CoV-2. The mNG SARS-CoV-2 derived from strain USA-WA1/20202 (WA1) was used as a template. The spike gene of WA1 was replaced with that of an individual variant. Amino acid mutations, deletions (del), and insertions (Ins) are indicated in reference to the USA-WA1/2020 spike. L: leader sequence; ORF: Open reading frame; NTD: N-terminal domain of S1; RBD: receptor binding domain; S: spike glycoprotein; S1: N-terminal Furin cleavage fragment of S; S2: C-terminal Furin cleavage fragment of S; E: envelope protein; M: membrane protein; N: nucleocapsid; UTR: untranslated region. BA.2 mutations of were depicted in gray; mutations unique to XBB.1.5 were shown in orange.

### Table S1. Twenty-eight human serum samples collected 15-32 days poster BA.5-bivalent-booster without prior SARS-CoV-2 infection

| Serum ID | Age (year) | Gender (F/M) | Race or Ethnicity | Serum collection time (days post-BA.5-bivalent booster) | Serum collection date | Last dose of mRNA vaccine before BA.5-bivalent booster | *$FFRNT_{50}$ | | | | | | | | | |
|---|---|---|---|---|---|---|---|---|---|---|---|---|---|---|---|---|
| | | | | | | | $USA-WA1/2020 | $BA.5-spike | XBB.1.5-spike | XBB.1.16-spike | XBB.2.3-spike | EG.5.1-spike | FL.1.5.1-spike | CH.1.1-spike | XBC.1.6-spike | BA.2.86-spike |
| 1 | 34 | F | Black | 15 | 9/30/2022 | Dose 3 | 5120 | 640 | 160 | 160 | 226 | 80 | 160 | 80 | 640 | 320 |
| 2 | 86 | M | White | 21 | 10/4/2002 | Dose 4 | 1280 | 28 | 10 | 10 | 10 | 10 | 10 | 10 | 10 | 10 |
| 3 | 31 | M | Asian | 20 | 10/6/2022 | Dose 2 | 5120 | 1280 | 160 | 160 | 80 | 40 | 40 | 40 | 320 | 160 |
| 4 | 61 | F | White | 15 | 10/11/2022 | Dose 4 | 5120 | 320 | 80 | 80 | 80 | 40 | 80 | 20 | 226 | 160 |
| 5 | 58 | F | Black | 14 | 10/11/2022 | Dose 3 | 7241 | 1810 | 160 | 113 | 160 | 160 | 160 | 80 | 1280 | 226 |
| 6 | 69 | M | White | 22 | 10/11/2022 | Dose 4 | 2560 | 80 | 10 | 20 | 10 | 10 | 20 | 10 | 80 | 28 |
| 7 | 67 | F | Asian | 21 | 10/12/2022 | Dose 4 | 20480 | 2560 | 80 | 57 | 160 | 80 | 113 | 20 | 453 | 160 |
| 8 | 77 | M | Asian | 27 | 10/13/2022 | Dose 4 | 2560 | 226 | 40 | 40 | 80 | 40 | 40 | 10 | 160 | 40 |
| 9 | 39 | F | White | 15 | 10/13/2022 | Dose 3 | 5120 | 320 | 20 | 14 | 20 | 10 | 10 | 10 | 40 | 10 |
| 10 | 73 | M | White | 24 | 10/13/2022 | Dose 4 | 1810 | 160 | 28 | 40 | 20 | 20 | 40 | 28 | 80 | 80 |
| 11 | 83 | F | White | 17 | 10/13/2022 | Dose 4 | 1280 | 40 | 14 | 14 | 20 | 20 | 10 | 10 | 40 | 20 |
| 12 | 79 | F | White | 26 | 10/16/2022 | Dose 3 | 3620 | 320 | 57 | 80 | 80 | 40 | 40 | 80 | 320 | 113 |
| 13 | 35 | F | Asian | 29 | 10/21/2022 | Dose 3 | 5120 | 160 | 40 | 40 | 40 | 20 | 20 | 20 | 320 | 80 |
| 14 | 73 | M | White | 26 | 10/17/2022 | Dose 3 | 7241 | 320 | 57 | 40 | 80 | 40 | 40 | 20 | 226 | 160 |
| 15 | 76 | M | White | 32 | 10/17/2022 | Dose 4 | 7241 | 1280 | 226 | 226 | 160 | 40 | 80 | 80 | 640 | 320 |
| 16 | 71 | M | White | 28 | 10/18/2022 | Dose 4 | 1280 | 40 | 10 | 14 | 10 | 10 | 10 | 10 | 20 | 20 |
| 17 | 22 | M | Hispanic | 19 | 10/18/2022 | Dose 3 | 10240 | 2560 | 160 | 80 | 160 | 160 | 80 | 80 | 1280 | 226 |
| 18 | 61 | F | White | 30 | 10/19/2022 | Dose 4 | 640 | 80 | 20 | 10 | 10 | 10 | 10 | 10 | 14 | 10 |
| 19 | 56 | M | White | 21 | 10/19/2022 | Dose 3 | 10240 | 640 | 80 | 113 | 160 | 40 | 40 | 40 | 320 | 113 |
| 20 | 66 | F | White | 26 | 10/19/2022 | Dose 4 | 3620 | 320 | 57 | 40 | 57 | 28 | 20 | 20 | 226 | 57 |
| 21 | 76 | F | White | 30 | 10/20/2022 | Dose 4 | 3620 | 226 | 10 | 10 | 14 | 10 | 10 | 10 | 80 | 28 |
| 22 | 61 | F | Asian | 31 | 10/20/2022 | Dose 3 | 10240 | 5120 | 453 | 320 | 320 | 320 | 226 | 160 | 5120 | 1280 |
| 23 | 77 | F | Black | 31 | 10/20/2022 | Dose 4 | 5120 | 80 | 10 | 10 | 20 | 10 | 10 | 10 | 40 | 40 |
| 24 | 59 | F | unknown | 28 | 10/21/2022 | Dose 4 | 2560 | 320 | 40 | 40 | 57 | 28 | 20 | 28 | 160 | 80 |
| 25 | 71 | M | Hispanic | 22 | 10/21/2022 | Dose 4 | 1280 | 160 | 20 | 20 | 40 | 20 | 14 | 10 | 80 | 28 |
| 26 | 70 | F | White | 22 | 10/21/2022 | Dose 3 | 2560 | 226 | 20 | 20 | 20 | 10 | 20 | 10 | 80 | 20 |
| 27 | 79 | F | White | 25 | 10/22/2022 | Dose 4 | 1280 | 113 | 10 | 10 | 20 | 10 | 10 | 10 | 28 | 20 |
| 28 | 79 | M | White | 18 | 10/22/2022 | Dose 4 | 2560 | 160 | 80 | 57 | 80 | 80 | 80 | 80 | 160 | 160 |
| Median | 70 | - | - | 23 | - | - | - | - | - | - | - | - | - | - | - | - |
| #GMT | - | - | - | - | - | - | 3576 | 290 | 42 | 39 | 48 | 29 | 31 | 23 | 152 | 67 |
| †95% CI | - | - | - | - | - | - | 2608-4903 | 173-485 | 27-64 | 26-58 | 31-72 | 20-42 | 21-45 | 16-33 | 87-267 | 42-108 |

*Individual $FFRNT_{50}$ value is the geometric mean of duplicate FFRNT results.
$Data was published previously[10].
#Geometric mean neutralizing titers (GMT).
†95% confidence interval (95% CI) for the GMT.

### Table S2. Twenty human serum samples collected 14-31 days poster BA.5-bivalent-booster with prior SARS-CoV-2 infection

| Serum ID | Age (year) | Gender (F/M) | Race or Ethnicity | Serum collection time (days post-BA.5-bivalent booster) | Serum collection date | Last dose of mRNA vaccine before BA.5-bivalent booster | *$FFRNT_{50}$ | | | | | | | | | |
|---|---|---|---|---|---|---|---|---|---|---|---|---|---|---|---|---|
| | | | | | | | [$]USA-WA1/2020 | [$]BA.5-spike | XBB.1.5-spike | XBB.1.16-spike | XBB.2.3-spike | EG.5.1-spike | FL.1.5.1-spike | CH.1.1-spike | XBC.1.6-spike | BA.2.86-spike |
| 1 | 19 | F | White | 19 | 10/4/2022 | Dose 3 | 10240 | 1280 | 160 | 80 | 80 | 80 | 160 | 40 | 640 | 226 |
| 2 | 69 | F | White | 15 | 10/6/2022 | Dose 4 | 10240 | 5120 | 453 | 320 | 320 | 320 | 320 | 320 | 1810 | 640 |
| [&]3 | 46 | F | White | 18 | 10/11/2022 | Dose 2 | 5120 | 2560 | 226 | 160 | 160 | 80 | 80 | 80 | 1810 | 320 |
| 4 | 80 | F | Black | 22 | 10/11/2022 | Dose 3 | 10240 | 7241 | 640 | 320 | 453 | 640 | 905 | 320 | 7241 | 640 |
| 5 | 76 | F | Hispanic | 21 | 10/12/2022 | Dose 3 | 5120 | 1280 | 160 | 80 | 160 | 113 | 160 | 80 | 1810 | 640 |
| 6 | 67 | M | White | 15 | 10/12/2022 | Dose 4 | 5120 | 2560 | 40 | 28 | 28 | 28 | 80 | 40 | 640 | 160 |
| 7 | 52 | F | White | 30 | 10/13/2022 | Dose 4 | 5120 | 1280 | 320 | 160 | 160 | 160 | 320 | 80 | 640 | 640 |
| 8 | 48 | F | Asian | 24 | 10/13/2022 | Dose 3 | 14482 | 7241 | 640 | 320 | 640 | 453 | 453 | 226 | 3620 | 2560 |
| 9 | 58 | F | White | 28 | 10/13/2022 | Dose 2 | 5120 | 640 | 40 | 40 | 40 | 20 | 40 | 28 | 226 | 80 |
| 10 | 67 | F | Black | 25 | 10/14/2022 | Dose 2 | 5120 | 2560 | 226 | 160 | 226 | 113 | 320 | 160 | 2560 | 160 |
| 11 | 20 | F | Hispanic | 21 | 10/14/2022 | Dose 2 | 1810 | 320 | 28 | 20 | 20 | 20 | 40 | 10 | 160 | 113 |
| [$]12 | 75 | M | White | 21 | 10/14/2022 | Dose 3 | 20480 | 2560 | 640 | 453 | 640 | 320 | 320 | 320 | 1810 | 1280 |
| 13 | 64 | F | White | 19 | 10/4/2022 | Dose 4 | 7241 | 2560 | 453 | 320 | 453 | 226 | 160 | 160 | 1810 | 1280 |
| 14 | 90 | F | White | 22 | 10/18/2022 | Dose 3 | 10240 | 1280 | 320 | 160 | 320 | 160 | 160 | 40 | 640 | 160 |
| [$]15 | 39 | M | Asian | 31 | 10/19/2022 | Dose 3 | 10240 | 5120 | 160 | 113 | 160 | 80 | 57 | 80 | 1810 | 226 |
| [$]16 | 67 | M | White | 17 | 10/19/2022 | Dose 4 | 10240 | 2560 | 320 | 160 | 320 | 320 | 320 | 160 | 1280 | 640 |
| [$]17 | 68 | M | Hispanic | 14 | 10/19/2022 | Dose 3 | 5120 | 2560 | 160 | 80 | 113 | 113 | 113 | 40 | 640 | 80 |
| 18 | 51 | M | White | 28 | 10/21/2022 | Dose 3 | 7241 | 905 | 80 | 80 | 57 | 40 | 40 | 20 | 226 | 113 |
| 19 | 65 | M | Asian | 29 | 10/20/2022 | Dose 4 | 2560 | 453 | 20 | 20 | 20 | 20 | 20 | 10 | 80 | 20 |
| 20 | 64 | M | White | 16 | 10/21/2022 | Dose 3 | 2560 | 640 | 20 | 10 | 20 | 14 | 10 | 10 | 320 | 40 |
| Median | 64 | - | - | 21 | - | - | - | - | - | - | - | - | - | - | - | - |
| [#]GMT | - | - | - | - | - | - | 6526 | 1810 | 157 | 100 | 130 | 97 | 121 | 65 | 874 | 260 |
| [†]95% CI | - | - | - | - | - | - | 4913-8668 | 1196-2740 | 91-271 | 61-166 | 75-225 | 56-166 | 71-207 | 38-112 | 513-1491 | 145-465 |

*Individual $FFRNT_{50}$ value is the geometric mean of duplicate FFRNT results.
[$]Data was published previously[10].
[&]Reported as SARS-CoV-2 RT-PCR positive. Nucleocapsid antibody testing was not performed. The rest of the samples were tested nucleocapsid-antibody positive.
[#]Geometric mean neutralizing titers (GMT).
[†]95% confidence interval (95% CI) for the GMT.

### Table S3. Forty-two human serum samples collected 15-117 days after XBB.1.5-infection from individuals who had received 2-4 doses of COVID-19 parental mRNA vaccine

| Serum ID | Age (year) | Gender (F/M) | Race or Ethnicity | [$]Serum collection time (days post-XBB.1.5 PCR+) | Serum collection date | Last dose of mRNA vaccine before XBB.1.5-infection | *$FFRNT_{50}$ | | | | | | | | | |
|---|---|---|---|---|---|---|---|---|---|---|---|---|---|---|---|---|
| | | | | | | | USA-WA1/2020 | BA.5-spike | XBB.1.5-spike | XBB.1.16-spike | XBB.2.3-spike | EG.5.1-spike | FL.1.5.1-spike | CH.1.1-spike | XBC.1.6-spike | BA.2.86-spike |
| 1 | 42 | M | Black | 15 | 2/2/2023 | Dose 3 | 3620 | 5120 | 1280 | 640 | 1280 | 1280 | 1280 | 320 | 7241 | 3620 |
| 2 | 64 | M | White | 16 | 2/10/2023 | Dose 2 | 20480 | 10240 | 1810 | 1280 | 2560 | 1810 | 2560 | 320 | 5120 | 1810 |
| 3 | 15 | F | Hispanic | 29 | 2/15/2023 | Dose 2 | 1280 | 905 | 160 | 160 | 160 | 160 | 160 | 80 | 640 | 320 |
| 4 | 27 | F | White | 31 | 2/24/2023 | Dose 3 | 5120 | 2560 | 640 | 640 | 640 | 640 | 640 | 320 | 1810 | 640 |
| 5 | 16 | M | Black | 39 | 2/27/2023 | Dose 2 | 2560 | 1810 | 905 | 640 | 1280 | 640 | 640 | 640 | 2560 | 1280 |
| 6 | 38 | F | White | 19 | 3/3/2023 | Dose 2 | 640 | 320 | 80 | 160 | 113 | 160 | 80 | 28 | 320 | 113 |
| 7 | 21 | F | White | 46 | 3/6/2023 | Dose 3 | 2560 | 320 | 80 | 40 | 80 | 40 | 40 | 14 | 160 | 113 |
| 8 | 46 | M | Hispanic | 35 | 3/6/2023 | Dose 2 | 905 | 640 | 226 | 320 | 320 | 453 | 320 | 113 | 905 | 640 |
| 9 | 40 | F | Hispanic | 39 | 3/11/2023 | Dose 2 | 640 | 320 | 160 | 80 | 160 | 160 | 226 | 40 | 640 | 905 |
| 10 | 56 | F | Black | 49 | 3/20/2023 | Dose 3 | 1280 | 905 | 80 | 160 | 160 | 80 | 160 | 80 | 640 | 320 |
| 11 | 42 | F | White | 25 | 3/22/2023 | Dose 2 | 640 | 320 | 80 | 80 | 160 | 80 | 113 | 40 | 320 | 320 |
| 12 | 21 | M | White | 40 | 3/22/2023 | Dose 2 | 640 | 320 | 80 | 80 | 160 | 113 | 80 | 40 | 453 | 160 |
| 13 | 36 | F | White | 19 | 3/22/2023 | Dose 3 | 1280 | 453 | 160 | 160 | 226 | 320 | 320 | 80 | 640 | 320 |
| 14 | 48 | F | White | 22 | 3/28/2023 | Dose 2 | 1280 | 905 | 226 | 320 | 320 | 640 | 905 | 160 | 2560 | 640 |
| 15 | 30 | F | White | 54 | 3/28/2023 | Dose 2 | 2560 | 640 | 80 | 160 | 160 | 80 | 113 | 80 | 453 | 226 |
| 16 | 61 | F | Hispanic | 82 | 4/18/2023 | Dose 3 | 320 | 160 | 40 | 40 | 40 | 57 | 57 | 20 | 453 | 40 |
| 17 | 25 | F | Hispanic | 42 | 4/18/2023 | Dose 3 | 5120 | 5120 | 1280 | 1280 | 3620 | 2560 | 2560 | 320 | 10240 | 5120 |
| 18 | 34 | F | Black | 45 | 4/20/2023 | Dose 2 | 640 | 320 | 80 | 113 | 160 | 160 | 160 | 40 | 453 | 160 |
| 19 | 53 | F | Hispanic | 73 | 4/27/2023 | Dose 2 | 1280 | 640 | 160 | 160 | 320 | 320 | 320 | 80 | 2560 | 640 |
| 20 | 40 | F | Hispanic | 92 | 5/4/2023 | Dose 2 | 320 | 160 | 40 | 40 | 80 | 80 | 80 | 20 | 320 | 113 |
| 21 | 26 | F | Hispanic | 48 | 5/4/2023 | Dose 2 | 905 | 640 | 80 | 80 | 113 | 80 | 113 | 40 | 640 | 160 |
| 22 | 27 | F | Hispanic | 37 | 5/4/2023 | Dose 2 | 5120 | 2560 | 640 | 640 | 1280 | 905 | 1280 | 320 | 5120 | 1280 |
| 23 | 71 | M | Asian | 31 | 5/4/2023 | Dose 2 | 1280 | 1280 | 453 | 453 | 640 | 640 | 640 | 160 | 1810 | 1280 |
| 24 | 56 | F | Hispanic | 49 | 5/4/2023 | Dose 3 | 1280 | 640 | 80 | 80 | 80 | 80 | 80 | 57 | 640 | 160 |
| 25 | 47 | F | White | 64 | 5/5/2023 | Dose 3 | 640 | 320 | 57 | 80 | 80 | 57 | 57 | 40 | 320 | 160 |
| 26 | 34 | F | White | 86 | 5/4/2023 | Dose 3 | 640 | 226 | 160 | 80 | 226 | 160 | 160 | 80 | 320 | 160 |
| 27 | 35 | M | Asian | 72 | 5/5/2023 | Dose 3 | 1280 | 320 | 57 | 40 | 57 | 40 | 80 | 40 | 320 | 320 |
| 28 | 40 | F | Hispanic | 96 | 5/8/2023 | Dose 2 | 453 | 113 | 28 | 20 | 20 | 20 | 10 | 20 | 80 | 20 |
| 29 | 39 | M | White | 65 | 5/9/2023 | Dose 2 | 1280 | 320 | 80 | 80 | 113 | 113 | 80 | 20 | 160 | 40 |
| 30 | 59 | F | Black | 42 | 5/9/2023 | Dose 3 | 3620 | 320 | 160 | 113 | 160 | 160 | 80 | 40 | 320 | 160 |
| 31 | 71 | F | White | 41 | 5/10/2023 | Dose 2 | 7241 | 10240 | 2560 | 2560 | 3620 | 2560 | 5120 | 1280 | 14482 | 7241 |
| 32 | 47 | M | White | 63 | 5/12/2023 | Dose 3 | 2560 | 905 | 320 | 640 | 640 | 320 | 453 | 160 | 905 | 320 |
| 33 | 27 | F | Hispanic | 52 | 5/15/2023 | Dose 2 | 2560 | 640 | 320 | 320 | 320 | 160 | 226 | 113 | 640 | 453 |
| 34 | 56 | F | Hispanic | 81 | 5/15/2023 | Dose 2 | 2560 | 1280 | 226 | 320 | 320 | 320 | 160 | 160 | 640 | 160 |
| 35 | 46 | F | White | 69 | 5/15/2023 | Dose 2 | 226 | 2560 | 640 | 453 | 640 | 640 | 320 | 905 | 5120 | 1280 |
| 36 | 59 | F | White | 38 | 5/16/2023 | Dose 3 | 2560 | 1280 | 640 | 640 | 640 | 320 | 320 | 320 | 1280 | 1280 |
| 37 | 30 | M | White | 117 | 5/16/2023 | Dose 3 | 1810 | 453 | 160 | 160 | 320 | 320 | 320 | 80 | 640 | 320 |
| 38 | 56 | F | White | 62 | 5/16/2023 | Dose 3 | 3620 | 1280 | 320 | 453 | 453 | 320 | 905 | 160 | 2560 | 905 |
| 39 | 40 | F | Hispanic | 79 | 5/18/2023 | Dose 4 | 1810 | 640 | 160 | 320 | 320 | 160 | 320 | 80 | 640 | 640 |
| 40 | 20 | M | Black | 35 | 5/22/2023 | Dose 2 | 10240 | 20480 | 2560 | 2560 | 2560 | 5120 | 3620 | 3620 | 20480 | 5120 |
| 41 | 28 | F | Asian | 75 | 5/22/2023 | Dose 3 | 1280 | 640 | 160 | 80 | 160 | 160 | 226 | 28 | 640 | 226 |
| 42 | 56 | M | Hispanic | 87 | 5/23/2023 | Dose 2 | 2560 | 5120 | 640 | 640 | 1280 | 1280 | 1280 | 640 | 10240 | 1280 |
| Median | 40 | - | - | 47 | - | - | - | - | - | - | - | - | - | - | - | - |
| [#]GMT | - | - | - | - | - | - | 1586 | 840 | 210 | 210 | 292 | 248 | 263 | 102 | 999 | 413 |
| [†]95% CI | - | - | - | - | - | - | 1176-2139 | 577-1223 | 146-303 | 146-303 | 200-427 | 168-366 | 174-397 | 69-152 | 665-1503 | 274-623 |

*Individual $FFRNT_{50}$ value is the geometric mean of duplicate FFRNT results.
[#]Geometric mean neutralizing titers (GMT).
[†]95% confidence interval (95% CI) for the GMT.

### Table S4. Nineteen human serum samples 17-103 days after XBB.1.5-infection from individuals who had received 2-4 doses of COVID-19 parental mRNA vaccine plus a BA.5-bivalent booster.

| Serum ID | Age (year) | Gender (F/M) | Race or Ethnicity | $Serum collection time (days post-XBB.1.5 PCR+) | Serum collection date | Last dose of mRNA vaccine before BA.5-bivalent booster | *$FFRNT_{50}$ | | | | | | | | | |
|---|---|---|---|---|---|---|---|---|---|---|---|---|---|---|---|---|
| | | | | | | | USA-WA1/2020 | BA.5-spike | XBB.1.5-spike | XBB.1.16-spike | XBB.2.3-spike | EG.5.1-spike | FL.1.5.1-spike | CH.1.1-spike | XBC.1.6-spike | BA.2.86-spike |
| 1 | 85 | F | White | 21 | 2/16/2023 | Dose 4 | 2560 | 453 | 160 | 160 | 160 | 113 | 80 | 40 | 320 | 160 |
| 2 | 81 | F | White | 22 | 2/2/2023 | Dose 4 | 1280 | 640 | 160 | 160 | 320 | 113 | 80 | 80 | 640 | 160 |
| 3 | 66 | F | Black | 20 | 2/3/2023 | Dose 3 | 10240 | 1280 | 453 | 640 | 1280 | 453 | 1280 | 320 | 5120 | 640 |
| 4 | 65 | M | Hispanic | 17 | 2/17/2023 | Dose 3 | 10240 | 2560 | 640 | 640 | 1280 | 640 | 1280 | 640 | 3620 | 2560 |
| 5 | 79 | F | Hispanic | 18 | 2/27/2023 | Dose 3 | 2560 | 453 | 160 | 160 | 320 | 160 | 320 | 40 | 640 | 320 |
| 6 | 68 | F | White | 53 | 3/13/2023 | Dose 3 | 2560 | 1280 | 320 | 320 | 640 | 320 | 320 | 160 | 1810 | 640 |
| 7 | 30 | F | Black | 52 | 3/15/2023 | Dose 2 | 1280 | 905 | 80 | 80 | 160 | 160 | 57 | 160 | 1280 | 160 |
| 8 | 79 | F | White | 50 | 3/17/2023 | Dose 4 | 5120 | 1280 | 113 | 160 | 160 | 80 | 80 | 80 | 1280 | 640 |
| 9 | 75 | F | White | 22 | 3/28/2023 | Dose 3 | 5120 | 2560 | 640 | 640 | 640 | 640 | 640 | 453 | 3620 | 640 |
| 10 | 71 | F | Black | 21 | 4/18/2023 | Dose 4 | 1810 | 1280 | 640 | 640 | 640 | 640 | 905 | 160 | 2560 | 905 |
| 11 | 72 | M | White | 23 | 4/18/2023 | Dose 4 | 1280 | 320 | 80 | 80 | 160 | 80 | 80 | 40 | 640 | 160 |
| 12 | 36 | F | Asian | 39 | 4/21/2023 | Dose 3 | 1280 | 160 | 40 | 57 | 80 | 80 | 57 | 28 | 226 | 80 |
| 13 | 63 | F | White | 56 | 4/26/2023 | Dose 3 | 453 | 113 | 20 | 20 | 28 | 40 | 28 | 10 | 57 | 20 |
| 14 | 68 | M | White | 66 | 4/28/2023 | Dose 4 | 5120 | 1280 | 320 | 320 | 640 | 320 | 453 | 160 | 2560 | 905 |
| 15 | 74 | M | White | 87 | 5/3/2023 | Dose 4 | 5120 | 640 | 160 | 226 | 453 | 226 | 160 | 160 | 1810 | 80 |
| 16 | 65 | M | White | 42 | 5/5/2023 | Dose 3 | 640 | 160 | 40 | 40 | 80 | 40 | 40 | 20 | 226 | 80 |
| 17 | 30 | F | Black | 103 | 5/5/2023 | Dose 2 | 905 | 640 | 80 | 40 | 113 | 113 | 40 | 57 | 640 | 113 |
| 18 | 29 | F | Hispanic | 42 | 5/16/2023 | Dose 3 | 5120 | 1810 | 320 | 320 | 320 | 320 | 320 | 160 | 1810 | 640 |
| 19 | 43 | F | Hispanic | 51 | 5/18/2023 | Dose 2 | 320 | 453 | 80 | 80 | 80 | 40 | 57 | 40 | 640 | 226 |
| Median | 68 | - | - | 42 | - | - | - | - | - | - | - | - | - | - | - | - |
| #GMT | - | - | - | - | - | - | 2173 | 689 | 154 | 160 | 252 | 163 | 160 | 88 | 956 | 267 |
| †95% CI | - | - | - | - | - | - | 1340-3523 | 443-1070 | 95-251 | 96-266 | 153-417 | 104-257 | 88-292 | 52-148 | 549-1665 | 151-470 |

*Individual $FFRNT_{50}$ value is the geometric mean of duplicate FFRNT results.
#Geometric mean neutralizing titers (GMT).
†95% confidence interval (95% CI) for the GMT.

### Table S5 *p* values calculated from Wilcoxon matched-pairs signed-rank test for group comparison of GMTs in Figure 1

| Figure Panels | P values from Wilcoxon matched-pairs signed-rank test |
|---|---|
| Figure 1A | USA-WA1/2020 versus all Omicron sublineage-spike: all <0.0001.<br>BA.5-spike versus other Omicron sublineage-spikes: all <0.0001.<br>XBB.1.5-spike versus XBB.1.16-, XBB.2.3-, EG.5.1-, FL.1.5.1-, CH.1.1-, XBC.1.6- and BA.2.86-spike: 0.157, 0.27, <0.0001, 0.0064, <0.0001, <0.0001, <0.0001.<br>XBB.1.16-spike versus XBB.2.3-, EG.5.1-, FL.1.5.1-, CH.1.1-, XBC.1.6- and BA.2.86-spike: 0.034, 0.11, 0.08, <0.0001, <0.0001, <0.0001.<br>XBB.2.3-spike versus EG.5.1-, FL.1.5.1-, CH.1.1-, XBC.1.6- and BA.2.86-spike: <0.0001, <0.0001, <0.0001, <0.0001, <0.0008.<br>EG.5.1-spike versus FL.1.5.1-, CH.1.1-, XBC.1.6- and BA.2.86-spike: 0.51, 0.06, <0.0001, <0.0001.<br>FL.1.5.1-spike versus CH.1.1-, XBC.1.6- and BA.2.86-spike: 0.0098, <0.0001, <0.0001.<br>CH.1.1-spike versus XBC.1.6- and BA.2.86-spike: both <0.0001.<br>XBC.1.6-spike versus BA.2.86-spike: <0.0001. |
| Figure 1B | USA-WA1/2020 versus all Omicron sublineage-spikes: all <0.0001.<br>BA.5-spike versus other Omicron sublineage-spikes: all <0.0001.<br>XBB.1.5-spike versus XBB.1.16-, XBB.2.3-, EG.5.1-, FL.1.5.1-, CH.1.1-, XBC.1.6- and BA.2.86-spike: <0.0001, 0.0039, <0.0001, 0.007, <0.0001, <0.0001, 0.0046.<br>XBB.1.16-spike versus XBB.2.3-, EG.5.1-, FL.1.5.1-, CH.1.1-, XBC.1.6- and BA.2.86-spike: 0.0015, 0.96, 0.20, 0.007, <0.0001, <0.0001.<br>XBB.2.3-spike versus EG.5.1-, FL.1.5.1-, CH.1.1-, XBC.1.6- and BA.2.86-spike: 0.018, 0.57, <0.0001, <0.0001, 0.0009.<br>EG.5.1-spike versus FL.1.5.1-, CH.1.1-, XBC.1.6- and BA.2.86-spike: 0.064, 0.0049, <0.0001, <0.0001.<br>FL.1.5.1-spike versus CH.1.1-, XBC.1.6- and BA.2.86-spike: 0.0004, <0.0001, 0.0032.<br>CH.1.1-spike versus XBC.1.6- and BA.2.86-spike: both <0.0001.<br>XBC.1.6-spike versus BA.2.86-spike: <0.0001. |
| Figure 1C | USA-WA1/2020 versus XBC.1.6- and other Omicron sublineage-spikes: 0.15, others <0.0001.<br>BA.5-spike versus XBC.1.6- and other Omicron sublineage-spikes: 0.018, others <0.0001.<br>XBB.1.5-spike versus XBB.1.16-, XBB.2.3-, EG.5.1-, FL.1.5.1-, CH.1.1-, XBC.1.6- and BA.2.86-spike: 0.91, <0.0001, 0.014, 0.0033, <0.0001, <0.0001, <0.0001.<br>XBB.1.16-spike versus XBB.2.3-, EG.5.1-, FL.1.5.1-, CH.1.1-, XBC.1.6- and BA.2.86-spike: <0.0001, 0.0165, 0.006, <0.0001, <0.0001, <0.0001.<br>XBB.2.3-spike versus EG.5.1-, FL.1.5.1-, CH.1.1-, XBC.1.6- and BA.2.86-spike: 0.019, 0.33, <0.0001, <0.0001, 0.0003.<br>EG.5.1-spike versus FL.1.5.1-, CH.1.1-, XBC.1.6- and BA.2.86-spike: 0.21, <0.0001, <0.0001, <0.0001.<br>FL.1.5.1-spike versus CH.1.1-, XBC.1.6- and BA.2.86-spike: all <0.0001.<br>CH.1.1-spike versus XBC.1.6- and BA.2.86-spike: both <0.0001.<br>XBC.1.6-spike versus BA.2.86-spike: <0.0001. |
| Figure 1D | USA-WA1/2020 versus XBC.1.6- and other Omicron sublineage-spikes: 0.003, others <0.0001.<br>BA.5-spike versus XBC.1.6- and other Omicron sublineage-spikes: 0.0005, others <0.0001.<br>XBB.1.5-spike versus XBB.1.16-, XBB.2.3-, EG.5.1-, FL.1.5.1-, CH.1.1-, XBC.1.6- and BA.2.86-spike: 0.19, 0.0001, 0.8, 0.4, 0.0004, <0.0001, 0.0006.<br>XBB.1.16-spike versus XBB.2.3-, EG.5.1-, FL.1.5.1-, CH.1.1-, XBC.1.6- and BA.2.86-spike: 0.0002, 0.52, 0.37, 0.0004, <0.0001, 0.002.<br>XBB.2.3-spike versus EG.5.1-, FL.1.5.1-, CH.1.1-, XBC.1.6- and BA.2.86-spike: 0.0005, 0.012, <0.0001, <0.0001, 0.477.<br>EG.5.1-spike versus FL.1.5.1-, CH.1.1-, XBC.1.6- and BA.2.86-spike: 0.44, <0.0001, <0.0001, 0.0012.<br>FL.1.5.1-spike versus CH.1.1-, XBC.1.6- and BA.2.86-spike: 0.0012, <0.0001, 0.016.<br>CH.1.1-spike versus XBC.1.6- and BA.2.86-spike: both <0.0001.<br>XBC.1.6-spike versus BA.2.86-spike: <0.0001. |
